# RNA-based communication in heterogeneous populations of cell mimics

**DOI:** 10.1101/2025.09.03.673985

**Authors:** François-Xavier Lehr, Imre Banlaki, Eric Bumüller, Thibault Mercier, Henrike Niederholtmeyer

## Abstract

RNA regulators offer a promising path for building complex, orthogonal circuits due to their low resource demands and design flexibility. In this study we explore their potential as signaling molecules in communication between synthetic cells. Specifically, we engineer populations of heterogenetic porous polymer cell mimics to produce, emit and receive two types of small synthetic RNA regulators. These RNAs are required to activate reporter expression at both the levels of transcription and translation. We distribute this AND gate circuit in receiver and two types of sender cell mimics to compare the distributed logic computation to the behavior of the circuit in well-mixed, bulk cell-free expression reactions. Analyzing different densities and spatial arrangements of senders and receivers, we reveal spatiotemporal gradients in RNA signals and identify configurations that increase specific activation. With small regulatory RNAs, the engineering toolbox for communication between synthetic cells expands to include a programmable class of signaling molecules. The rapid turnover of RNA suggests applications in establishing dynamic signaling gradients in communities of synthetic cells.

**Graphical Abstract:** 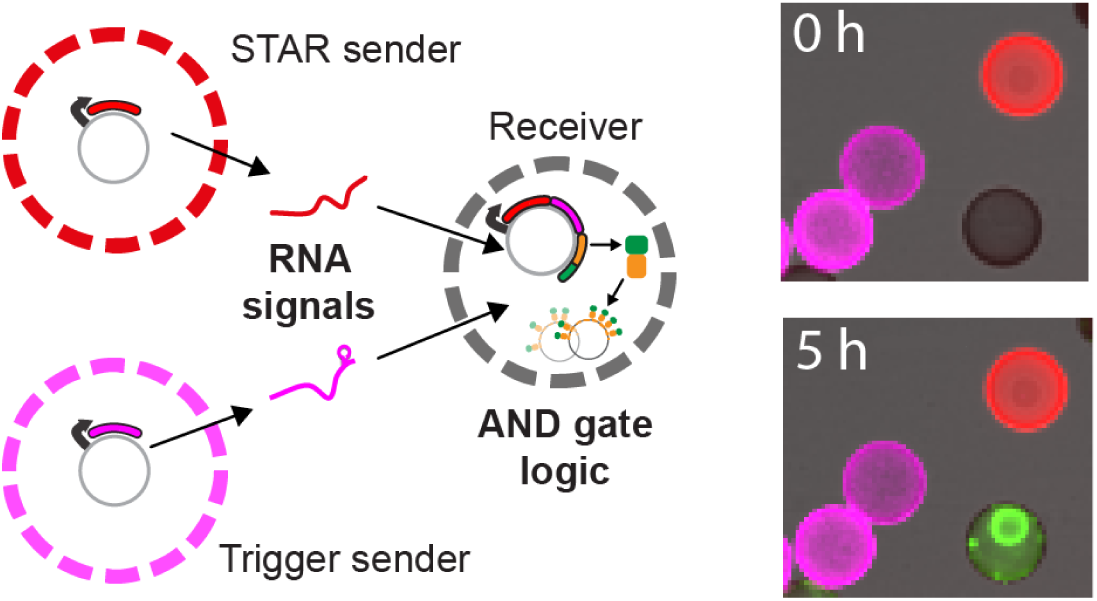

## Introduction

Emulating cellular functions by constructing synthetic cells from non-living matter allows us to learn more about the essential features of life. ^1^ Bottom up assembly of synthetic cells has led to fundamental insights in cell biology. ^2^ Control over important parameters in simplified mimics of living cells enables systematic analyses that are difficult to perform in living, natural systems. Synthetic, cell-like compartments can make use of biological materials and functions from different organisms, like whole cell extracts, recombinant DNA and proteins, as well as non-natural materials, so many practical applications can be envisioned. Synthetic cells could be genetically programmable, soft microrobots ^3–5^ that sense, and react to environmental signals, for example as smart drug delivery agents, to interact with living cells in regenerative medicine, in bioremediation and in novel biomanufacturing routes. However, the biosynthetic capabilities of synthetic cells remain limited, constraining the development of sensing and actuation functions.^6–8^

Intercellular communication is essential in multicellular and many single celled organisms to coordinate collective behaviors and regulate responses of specialized cells. Communication between non-living, synthetic cells could help with some of their limitations, for instance to optimize the use of limited resources by dividing labor between specialized populations of synthetic cells. Releasing and processing signals allows cells to influence and respond to their local environment, a prerequisite for accurate responses in practical applications and the formation of patterns in synthetic tissues and materials composed of synthetic cells. Recognizing the importance of intercellular communication, the engineering of new signaling routes between synthetic cells has recently received a lot of attention. ^9–12^ Besides expanding the communication toolbox for synthetic cells, this research improves our understanding of natural systems, as the programmability of minimal systems enables systematic studies on the influence of spatial arrangements between senders and receivers. ^13–16^ An understanding of signaling distances and how signals from different sources are integrated is essential to engineer predictable responses.

The type of compartment, its permeability and its biochemical functionalization influence which molecular signals a synthetic cell can produce, release and sense. The two main approaches in synthetic cell communication are microfluidic chambers and free, capsule-like compartments. The advantage of microfluidic chambers is their high precision in spatial connectivity and experimental control. ^16,17^ On the other hand, free-floating, capsule-like compartments enable the use of a wider range of materials for interface design, higher compartment numbers and flexible arrangements in continuous one- to three-dimensional spaces. ^18–20^ For capsule-like compartments, intercellular communication depends on membrane permeability. Membranes of phospholipid vesicles are permeable to small, apolar molecules such as acyl homoserine lactones from bacterial quorum sensing, ^21–23^ which can be synthesized and converted to a gene expression response by genetically encoded synthases and transcription factors. To expand the range of signals, alpha-hemolysin pores within phospholipid membranes allow the release and uptake of small polar molecules that serve as substrates or inducers for a response in the synthetic receiver cells. ^14,24–26^ Contact-dependent and light-mediated signaling are other communication mechanisms that are compatible with lipid membranes. ^18,19,27–29^ Semipermeable polymeric capsules are more stable alternatives to compartments with lipid membranes and support even wider ranges of diffusive signaling molecules. For example, we have recently developed cell mimics with porous polymethacrylate membranes that can synthesize and release protein signals. ^30,31^ Proteinosomes with crosslinked protein membranes are permeable to oligonucleotide signals in DNA strand displacement reactions that perform computations in receiver compartments. ^13,27,32,33^

Nucleic acids are exciting as signaling molecules because their interactions with target nucleic acids can be easily engineered to design orthogonal regulators. RNA is particularly interesting because it regulates and encodes protein synthesis, and proteins are required for most advanced functions of synthetic cells. However, RNA-signals in synthetic cell communication, especially to regulate coupled transcription and translation reactions, remain mostly unexplored. So far, diffusive RNA signals traveling between proteinosomes regulated CRISPR/Cas computations in transcription-only synthetic circuits, ^33^ and even more recently, the Adamala lab used cell-penetrating peptides to send an mRNA from one synthetic cell population to another. ^34^

In this study, we use porous polymer cell mimics that we program with genetic circuits. We previously demonstrated communication through protein signals that influenced gene expression in neighboring cell mimics. ^30^ Here, we aimed to establish RNA-based communication to regulate protein synthesis in heterogeneous populations of sender and receiver cell mimics. As signals, we chose two small RNA regulators: a Small Transcription Activating RNA (STAR) as a transcriptional activator, ^35^ and a trigger RNA activating a Toehold switch at the translation level. ^36^ These two regulators were previously assembled into an AND gate that controlled protein synthesis in bulk cell-free expression reactions. ^37^ AND gate logic allows us to separate the two activating signals into two distinct sender populations and to investigate how activation of receiver cell mimics depends on the densities and spatial arrangements of the communicating cell mimics.

We demonstrate successful RNA-based communication and spatially distributed logic computation in genetically heterogeneous cell mimic communities composed of two sender populations that activate receivers harboring the AND gate. We find that densities and localization of the communicating cell mimics with respect to each other lead to differential AND gate activation in individual receivers. We identify leakiness and low signal to noise ratios as challenges in the distributed RNA communication network. Fold-activation of receivers improves when both activating RNAs emit from a nearby source, around which a spatiotemporal gradient forms. With small regulatory RNAs, the engineering toolbox for synthetic cells expands to include a highly programmable class of signaling molecules.

## Results and discussion

### Design of the RNA communication circuit for cell mimics

To engineer RNA-based communication between cell mimics, we chose an AND gate consisting of two small, synthetic RNA activators. ^37^ To activate protein production, the AND gate requires the presence of both RNA activators: A short RNA that activates transcription (STAR) ^35^ and a short trigger RNA that activates a toehold switch to allow translation. ^36^ This multilevel circuit comprises three DNA constructs: two activator constructs encoding the STAR and the trigger RNAs, as well as the AND gate construct. The AND gate is a reporter construct, regulated by the target region for the STAR and by a trigger-activated toehold switch upstream of the gene for a fluorescent reporter protein fused with TetR (TetR-sfGFP) (**Fig. 1A**).The fusion with TetR is important for capture of the reporter protein in cell mimics, which will be explained later. To activate production of the reporter, the STAR antisense RNA disrupts the formation of an intrinsic terminator. In the absence of the STAR, rho-independent termination ends transcription early, before reaching the toehold switch and coding region. The trigger-activated toehold switch regulates gene expression at the translation level. Without the trigger, a stem loop structure sequesters the region around the ribosomal binding site (RBS), preventing translation. Binding of the trigger allows for favorable refolding of the toehold stem, rendering the RBS accessible to the ribosome and enabling translation of the transcript. Consequently, the reporter gene is only transcribed and translated in the presence of both small RNA molecules (**Fig. 1A**).

**Figure 1.**
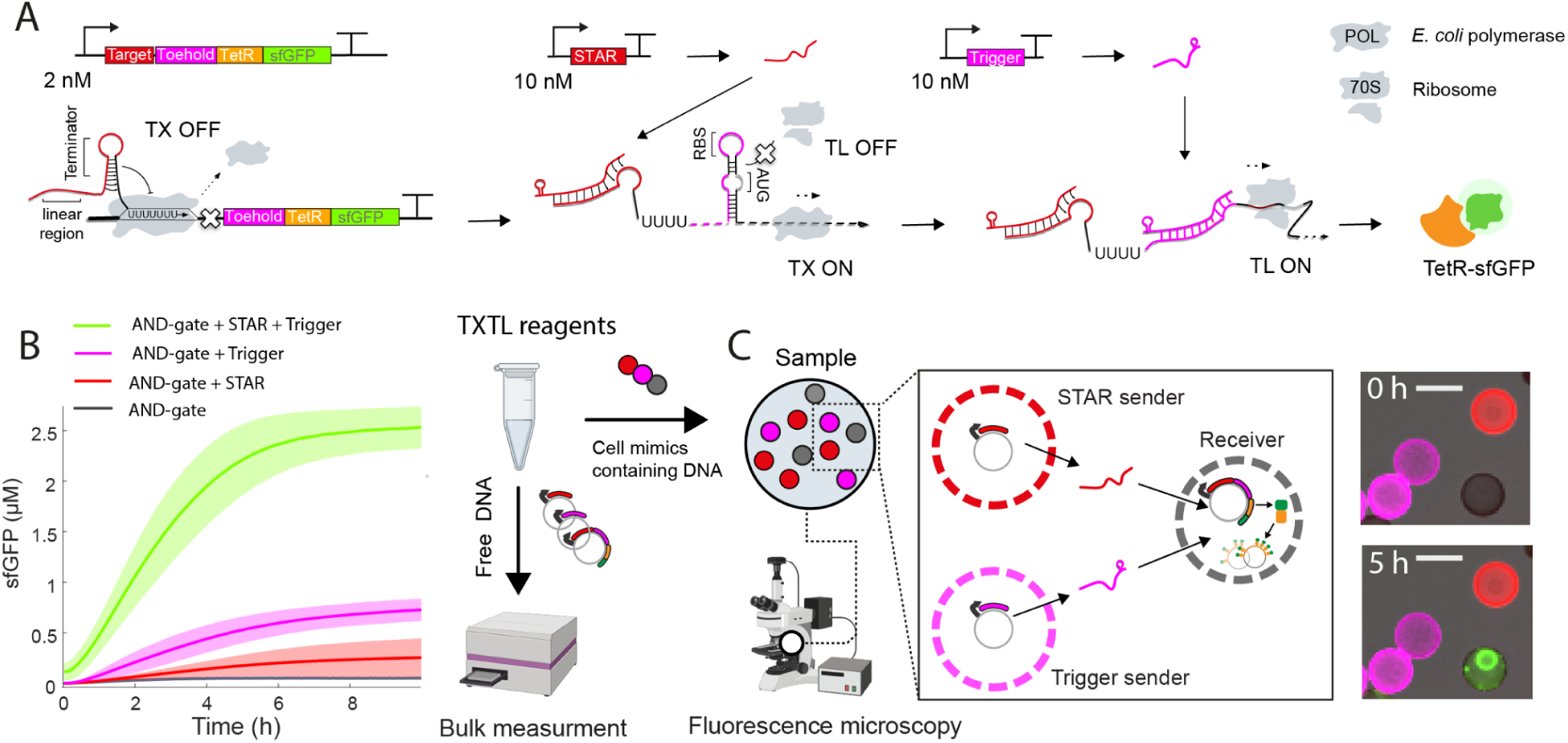
Design and testing of a RNA communication circuit for cell mimics. A) Scheme of the AND-gate circuit consisting of three separate plasmids. Labels indicate concentrations in bulk reactions. B) Kinetics of bulk reactions testing the AND gate circuit. Bold lines represent the mean values, and shaded areas indicate the standard deviation calculated from three biological replicates. C) Scheme of RNA-based communication in cell mimics. Circuit plasmids are distributed into three distinct cell mimic populations that can be differentiated by their polymer shell fluorescence. Microscope images show a region of a cell mimic experiment right after immersing cell mimics in TXTL reagents (top) and at the end of the reaction (bottom), with STAR (red) and trigger senders (magenta), and TetR-sfGFP signal (green) concentrated within the condensed clay hydrogel nucleus in a receiver. Scale bars: 70 µm.

First, we tested the circuit in bulk, cell-free transcription and translation reactions (TXTL) that contained free DNA (**Fig. 1B**). We used high concentrations of STAR and trigger DNA templates (10 nM each) to ensure maximal activation of the AND-gate. As expected, reactions containing both activator constructs to synthesize STAR and trigger RNA produced the highest sfGFP signals, while the lowest signal was detected in their absence. In controls with only one activator, the trigger activator alone produced a higher leakage compared to the STAR activator alone, which is consistent with previous results. ^37^ In bulk reactions with free DNA, this resulted in a 3.5-fold difference between the full AND gate with both activators and the addition of trigger alone, which we deemed sufficient for further experiments. The increased leakiness of the AND gate circuit as compared to a previous implementation could be due to a different reporter coding sequence downstream of the toehold switch in our design, which may affect the stability of the stem loop structure that blocks access to the RBS. ^37^ However, we cannot rule out other reasons such as the high concentration of trigger plasmid used in our experiments compared to previously tested ranges (2-7 nM) and *E. coli* lysate batch effects on mRNA half-life and production rate.

After successful verification of the circuit in bulk TXTL reactions, we aimed to test if STAR and trigger RNA could serve as signaling molecules that diffusively travel between cell mimics and activate reporter gene expression in receivers. As cell mimics we use porous polymer compartments, in which we immobilize DNA templates in a clay-DNA hydrogel nucleus. To prepare DNA-loaded compartments, we incubate polymer shells in a suspension of synthetic clay and DNA, which diffuse through the porous polymer membrane (see Methods). Upon addition of ions, the clay mineral discs form a condensed hydrogel that tightly binds DNA molecules in cell mimics and is too large to exit the polymer cage. ^30,31^ The polymer membranes of the compartments are permeable for all components of a TXTL reaction, even ribosomes, allowing us to initiate expression of DNA templates immobilized within cell mimics by simply immersing them in TXTL reagents. We hypothesized that regulatory RNAs such as the STAR and trigger activators should be able to travel between neighboring cell mimics in this system. To test RNA-based communication, we prepared three distinct populations of cell mimics, each containing one of the circuit’s DNA templates: STAR senders, trigger senders and receivers (**Fig. 1C**). To distinguish the cell mimic populations from each other, we fluorescently labeled the polymer membranes of the senders. Trigger senders were labeled with CF-555 dye (depicted in magenta) and STAR senders with CF-633 dye (depicted in red). The polymer membranes of receivers remained unstained. Receiver cell mimics were loaded with the AND gate reporter plasmid and a tetO array plasmid, which has a large number of TetR repressor binding sites to capture the reporter fusion protein (TetR-sfGFP). This means that synthesized reporter protein is captured by its producing, and nearby, receiver cell mimics localizing the reporter signal to its region of synthesis. ^30^ We loaded the sender cell mimics with higher DNA amounts than receivers, while staying within the limits of DNA handling and clay-core binding capacity, because we found that senders needed to be loaded with high DNA concentrations to produce a detectable reporter output signal in receiver cell mimics.

In cell mimic communication experiments, we immerse cell mimics in TXTL reagents to initiate the reaction. For imaging, the cell mimic suspension is added on a microscopy dish and sandwiched with cover glass to prevent evaporation and to produce a round droplet with dispersed cell mimics (**Fig. 2A**). In this configuration, transcription is localized within cell mimics. When transcription is complete, RNA molecules can diffuse through the porous membrane, allowing regulatory RNAs to find their targets. Even though ribosomes are slightly enriched within the clay-DNA hydrogel, ^30^ where mRNA is produced, translation can take place within as well outside of cell mimics. If the reporter protein is produced, receiver cell mimics capture the TetR-sfGFP reporter and the clay-DNA hydrogel within them becomes visible (**Fig. 1C**).

**Figure 2.**
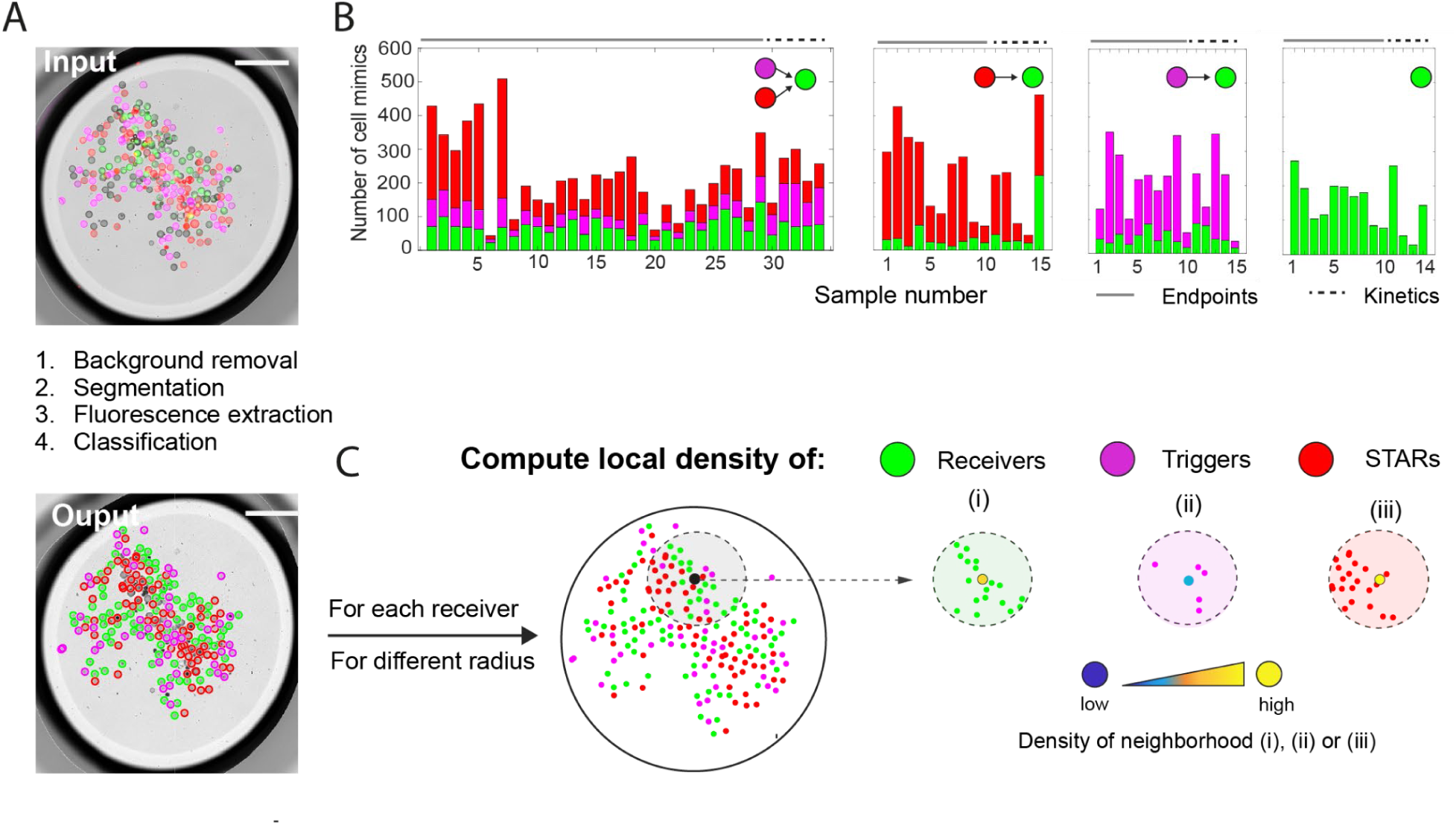
Image analysis pipeline and distribution of cell mimics in our experiments.(A) After background correction of the fluorescence channels, segmentation, and fluorescence extraction, the identified cell mimics are classified as receiver, trigger sender, or STAR sender in each sample. (B) Overview of the numbers of cell mimics per population type for all samples analyzed in this study, including endpoint (analysis in Fig. 3) and kinetic measurements (analysis in Fig. 4). (C) Local densities of receivers, STAR, and trigger senders are computed for each individual receiver. The example shows how the area used to compute the local density depends on the radius of the circle around the respective receiver. Scale bars: 700 µm

### Design and analysis of communication experiments

As both small RNAs are required to activate transcription and translation of the reporter gene located in receiver cell mimics, we wondered how the density and spatial arrangement of the three cell mimic populations with respect to each other influenced activation. In order to analyze communication via the STAR and the trigger RNA signals, and to answer our questions about the importance of densities and spatial arrangement of the communicating cell mimics, we took advantage of the random positioning of cell mimics when we prepared samples for microscopy. When pipetting and sandwiching a small volume of cell mimics in a microscopy dish, cell mimics of the different populations (STAR senders, trigger senders and receivers) settle at random positions within the TXTL droplet (we call this a sample) (**Fig. 2A**). We analyzed a large number of samples, in which we varied the amounts of the three populations of cell mimics while keeping the volume of external TXTL solution constant at 5 µl. We also included control experiments, in which we omitted one or both of the senders (**Fig. 2B**). Based on the kinetic data of our bulk experiments, we imaged most cell mimic samples after 5 h to get an endpoint measurement, but for a smaller number of experiments we performed time-lapse imaging to get kinetic data for the activation of all receivers within a sample.

In total, we acquired endpoint images for 60 samples, and performed time-lapse imaging for 19 samples. After processing the images, we segmented them and automatically classified cell mimics into the three populations based on their extracted fluorescence signatures (**Fig. 2A**). The numbers of cell mimics per sample varied, with some containing as few as 28 cell mimics and others exceeding 500 (**Fig. 2B**). While the number of senders were weakly or not correlated with the number of receivers in the experiments, STAR and trigger sender amounts were correlated because they were premixed during sample preparation (**Fig. S1**). In addition to quantifying the sender and receiver populations globally for each sample, we introduced an additional parameter to reflect the local neighborhood composition around each receiver. We calculated the local density of each population (STAR or trigger senders, and receivers) within a specified radius around each individual receiver (**Fig. 2C**). At the end of this analysis pipeline, each receiver cell mimic across all samples was assigned a sfGFP fluorescence value, a local density for each type of neighboring cell mimic populations, and its 2D coordinates. To test whether positional effects within the sample might influence activation, we assessed the correlation between the x- and y-coordinates of receivers and their sfGFP fluorescence. No significant correlation was observed (Pearson’s r < 0.1 for both x and y across all samples; **Fig. S2**), confirming that activation is not significantly influenced by x-y position in the sample.

### Global and local cell mimic densities influence activation of receivers

Receiver cell mimics have to integrate signals from two sources and, with about 17 minutes for a typical mRNA, ^38^ the lifetime of RNA in TXTL reactions is limited. We can therefore expect that activation of receiver cell mimics will depend on the density of each of the senders in a given experiment, and that activation of individual receivers may depend on their distance to senders. Visually, receiver fluorescence indeed increased with increasing numbers of sender cell mimics in the sample (**Fig. 3A**).

**Figure 3.**
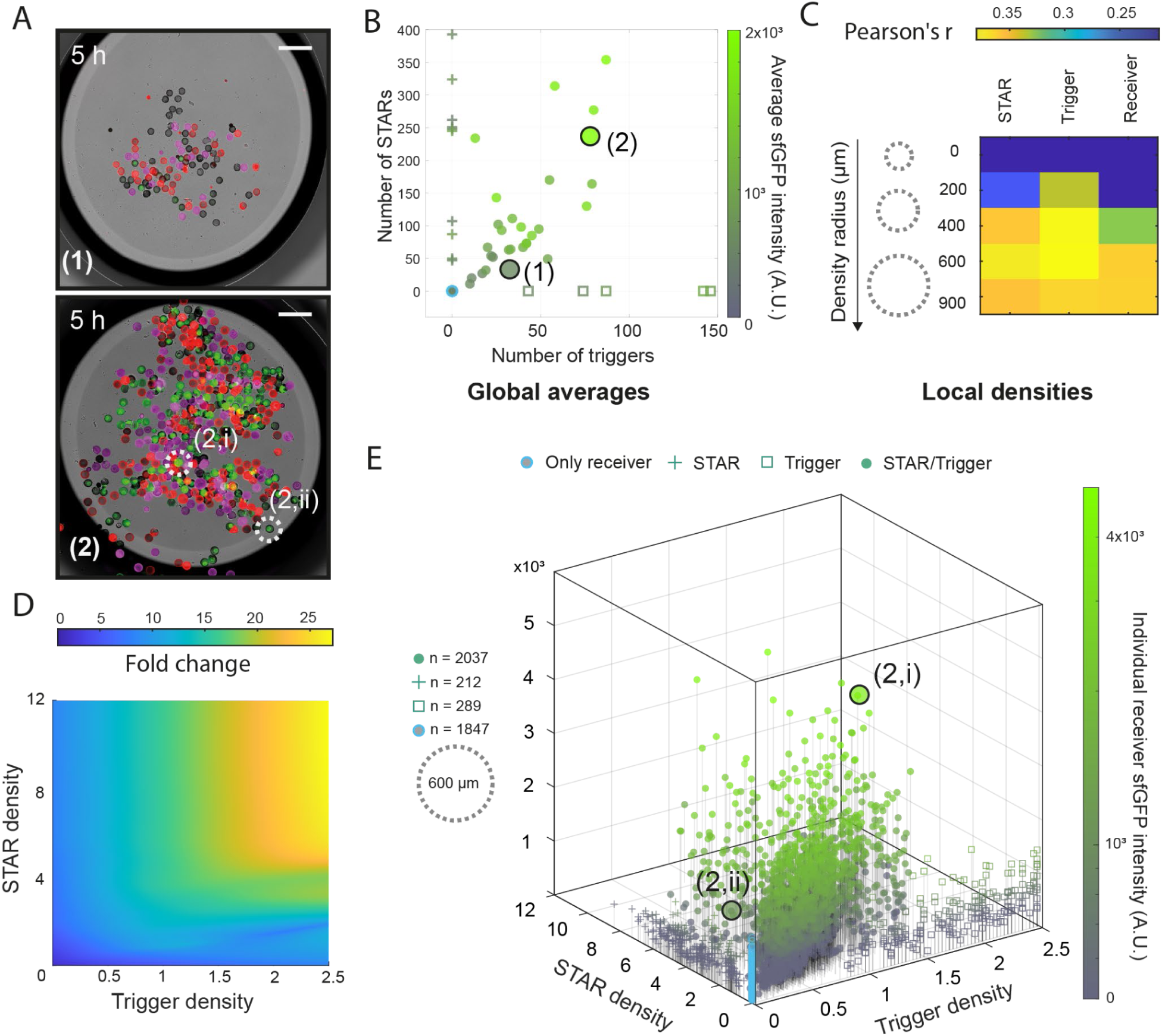
Verification of RNA-based communication and the AND-gate circuit in cell mimics using endpoint data. (A) Examples for experiments with a low (1) and a high (2) number of cell mimics in the sample, imaged after 5 hours. Merge of brightfield and fluorescence channels for sfGFP (green), STAR senders (red) and trigger senders (magenta). Scale bars: 400 µm. (B) Analysis of sfGFP endpoint fluorescence values averaged across all receivers in a sample. Samples with all three cell mimic types were compared to control experiments that omitted senders. Data points from the examples in (A) are indicated. (C) Pearson correlation coefficients (r) were calculated to assess the correlation between the measured sfGFP signals for receiver cell mimics and the density of the three cell mimic populations in a radius around a receiver cell mimic. The radius with the highest correlation coefficients (600 µm) was chosen for further analyses. (D) LOESS-smoothed surface showing fold change of individual receiver sfGFP fluorescence intensities as a function of trigger and STAR densities. Fold changes are computed relative to the averaged intensities of cell mimics in the absence of both senders. (E) Individual receiver sfGFP fluorescence intensities across all experiments, including controls that omitted senders, plotted against local sender densities in a radius of 600 µm around each receiver. Density values indicate the number of cell mimics within the radius around the receiver divided by the circle area. Data points from two receiver cell mimics in (A) sample (2) are indicated. Density values are the number of cell mimics per mm^2^.

Before detailed analyses of local environments, we decided to first look at receiver activation globally, to determine how the number of each of the three cell mimic types affects the average receiver activation in a given sample. For the 29 endpoint samples containing both sender types and receivers, a distinct increase in the average sfGFP signal was observed with a rising number of STAR or trigger senders (**Fig. 3B**). Notably, the sfGFP signal remained elevated with fewer trigger senders (as low as 12) but required a larger number of STAR senders (approximately 100) to be visible. This discrepancy in sender number requirements reflects the results from our own and previous bulk experiments with free DNA, where only 2 nM of trigger but 10 nM of STAR maximized the fold-change of the activated AND gate. ^37^ Comparing experiments containing up to 150 trigger cell mimics, control experiments with receivers and only one sender type, or no sender cell mimics at all, confirmed that both signals were required for the strongest activation of reporter gene expression in receivers (**Fig. 3B**). Consistent with our bulk experiments, we did observe some leaky expression in control experiments. We measured the lowest sfGFP signals in the absence of both senders. We observed a slight increase in sfGFP intensity when only STAR senders were present, regardless of the numbers of STAR senders (50-400). In experiments with only trigger senders, leaky expression was generally higher than in STAR sender only experiments, but remained lower than in experiments with both sender types (**Fig. 3B**). Leaky expression signals only increased for samples containing high numbers of trigger senders (>150) that went beyond the range of tested trigger sender numbers in experiments with both sender types. Beyond 150 trigger cell mimics in a sample, sfGFP signals increased with increasing trigger numbers, even in the absence STAR senders, indicating that some leakage occurs at the level of transcription termination (**Fig. S3**). The number of receiver cell mimics in a sample had little effect on sfGFP signals (**Fig. S1, S3, S7**).

Our analyses of global receiver activation verified RNA-based communication and AND-gate computation in cell mimics. However, the fold-changes in sfGFP signal averaged across the receiver cell mimics within a given sample were notably lower compared to bulk experiments. A possible reason for this is that our cell mimic experiments contained lower total DNA concentrations (approximately 0.36 nM of AND gate reporter plasmid for a sample with 100 receivers, or 2 nM of RNA activator plasmid for a sample with 100 senders). Locally however, close to individual cell mimics, DNA concentrations and activator synthesis rates are high. We hypothesized that activation may best be observed locally, while averaging across the spatially heterogeneous sample hides strong activation of receivers in local hot spots. In images, we indeed often observed high sfGFP signals in dense areas (**Fig. 3A**). To explore and quantify this, we calculated the local density of STARs, triggers, and receivers around each receiver for various radii (**Fig. 2C**). Using the local densities of cell mimics within different radii around a given receiver, we assessed the correlation between its sfGFP signal and these densities, first within each sample and then on average across all samples (**Fig. 3C**). The highest averaged correlation coefficients for all three cell mimic types were observed at a radius of 600 µm, with moderate and significant correlations for STAR and trigger densities (Pearson’s r = 0.40, p-values = 0.042 and 0.047 respectively) and a weaker, averaged correlation for receiver densities (Pearson’s r = 0.34, p-value = 0.065). Correlation coefficients varied widely across samples (up to 0.65, down to near zero), likely due to the broad distribution of cell mimic numbers in the samples. We find the highest correlations between sender density and receiver sfGFP signals in samples with intermediate numbers of STAR and trigger senders (**Fig. S4**). We hypothesize that samples with a high number of senders have lower correlation coefficients because there are fewer spatial configurations where a receiver contains a low amount of senders in its neighborhood. Conversely, samples with a very low number of senders suffer from the reverse effect, where fewer spatial configurations activate a receiver. Other factors that contribute to variability may be differences in DNA loading between individual cell mimics ^30^ (**Fig. S5**) and the random positioning of cell mimics in samples (**Fig. 3A**), which could lead to differences in convective flows due to slight drying and temperature gradients during sample incubation.

At a small radius of 200 µm, the correlation coefficients were low for both STAR and trigger densities, suggesting that an area of 0.12 mm^2^ is not big enough to achieve sufficient concentrations of STAR and trigger RNAs to activate the close-by receiver. The relatively higher correlation coefficient for the trigger senders at this small radius suggests that the trigger signal is more potent, requiring fewer senders, and acts more locally (**Fig. 3C**). We computationally predicted the 3D structures and diffusion coefficients of the signal RNAs. STAR is estimated to be more structured with a diffusion coefficient of 8.48*10^−7^ cm^2^/s, whereas the less folded trigger RNA is estimated to have a diffusion coefficient of 6.49*10^−7^ cm^2^/s (**Fig. S6**). The more localized activity of the trigger RNA may be due to its stronger effect at lower concentrations (**Fig. 3B**), as well as its lower predicted diffusion coefficient as compared to the STAR (**Fig. S6**).

Plotting the sfGFP endpoint values of individual receivers against their local densities of trigger and STAR senders within a radius of 600 µm, where the highest correlation coefficients were observed, confirmed that only receivers with both senders in the sample exhibit high sfGFP signals (**Fig. 3D, 3E, Movie 1**). Considering activation of individual receivers in areas with high local densities of both senders, we now observe similar dynamic ranges between ON and OFF states as in the bulk experiments of over 25-fold (**Fig. 3D**). The most active receivers were located in areas with a high density of STAR (>4 STAR senders/mm^2^) and an intermediate to high density of trigger senders (>0.5 trigger senders/mm^2^) (**Fig. 3E**). Mirroring the results from bulk experiments, ^37^ we find that a local concentration of STAR senders 5 to 10 times higher than that of trigger senders is required for optimal activation. Comparing individual sfGFP signals, we note that not all receivers in a similar local environment activate to the same extent. From a previous study ^30^ we know that cell mimics are not uniform in their ability to produce and bind the reporter protein, and we indeed observe differences in DNA loading between individual cell mimics (**Fig. S5**).

The distribution of single receiver fluorescence in control experiments provided deeper insights into the main source of AND-gate leakiness than the averaged data. In the absence of senders or with only STAR senders present, most receivers exhibited low fluorescence levels, regardless of STAR sender density (**Fig. S7**). This suggests that regulation at the level of translation is tight. The leaky expression we had observed in the averaged receiver fluorescence in trigger-only samples could be attributed mainly to receivers in high-density trigger sender areas (>2.5), suggesting that termination of transcription in the absence of STAR activators does not occur as reliably as regulation at the level of translation. Therefore, if the local concentration of trigger RNA is high, this leads to production of the reporter protein.

Apart from an inherent leakiness previously observed for STAR regulators, ^37,39,40^ we cannot exclude that mRNA interactions with the clay hydrogel inhibit or delay the formation of the terminator hairpin. The strong affinity of clay minerals for nucleic acids and its proximity during transcription makes interactions likely. ^41^ However, the distributed AND gate shows that *in situ* synthesized RNA is not irreversibly immobilized. Depending on size and secondary structures, RNAs may experience multiple transient binding events modifying their activity and diffusive mobility. Purposefully using transient binding interactions may be another tool to engineer RNA signal propagation in synthetic cell communities in the future.

### Activation kinetics reveal dynamic gradients of RNA signals

Because leaky expression, diffusion of RNA signals, as well as diffusion of reporter mRNA and protein within a sample may smoothen local effects and increase overall sfGFP signals with time, we analyzed the kinetics of activation in individual receiver cell mimics. To do so, we performed time-lapse imaging for a subset of experiments (**Movie 2**). In experiments containing both STAR and trigger senders together with receivers, we calculated the Pearson correlation coefficient between the local population densities and the fluorescence of individual receivers at each time point (every 15 minutes) (**Fig. 4A**). The highest correlation coefficients were observed between 1.5 h and 3.5 h (Pearson’s r = 0.43). The high correlation at the intermediate time points was due to a rapid increase in sfGFP intensities in receivers located in densely populated areas (**Fig. 4B and C**). Notably, after 3.5 h, the correlation between receiver fluorescence and densities of all cell mimic types in the local neighborhood began to decline. Despite the decrease in correlation, at 5 hours, corresponding to the endpoint values in our previous analyses (**Fig. 3**), we still observed significant Pearson correlation coefficients. The reduced correlation at 5 hours could be attributed to an accumulation of sfGFP signal in isolated receiver cell mimics located in less densely populated regions toward the end of the time-lapse experiments. Here, sfGFP signals initially increased very slowly, but often started to rise at about 4 h (**Fig. 4C**). In control experiments (**Fig. 4D**), where we omitted one type of sender, we observed a similar behavior as in these isolated receivers. Independent of STAR sender density, the kinetic traces largely remained flat, with a minor increase observed towards the end of the reaction (**Fig. 4D**). In samples containing only trigger senders, we again observed higher leakage, manifesting as earlier sfGFP production correlating with increased trigger density around the receivers. In areas with lower trigger densities, leaky expression was low in the initial phase of the reaction and only increased during the later phase of the reaction (**Fig. 4D**). The receiver density exhibited the weakest correlation with receiver fluorescence, and did not display any consistent trend in the kinetics of sfGFP signal accumulation (**Fig. 4A, Fig. S8**).

**Figure 4.**
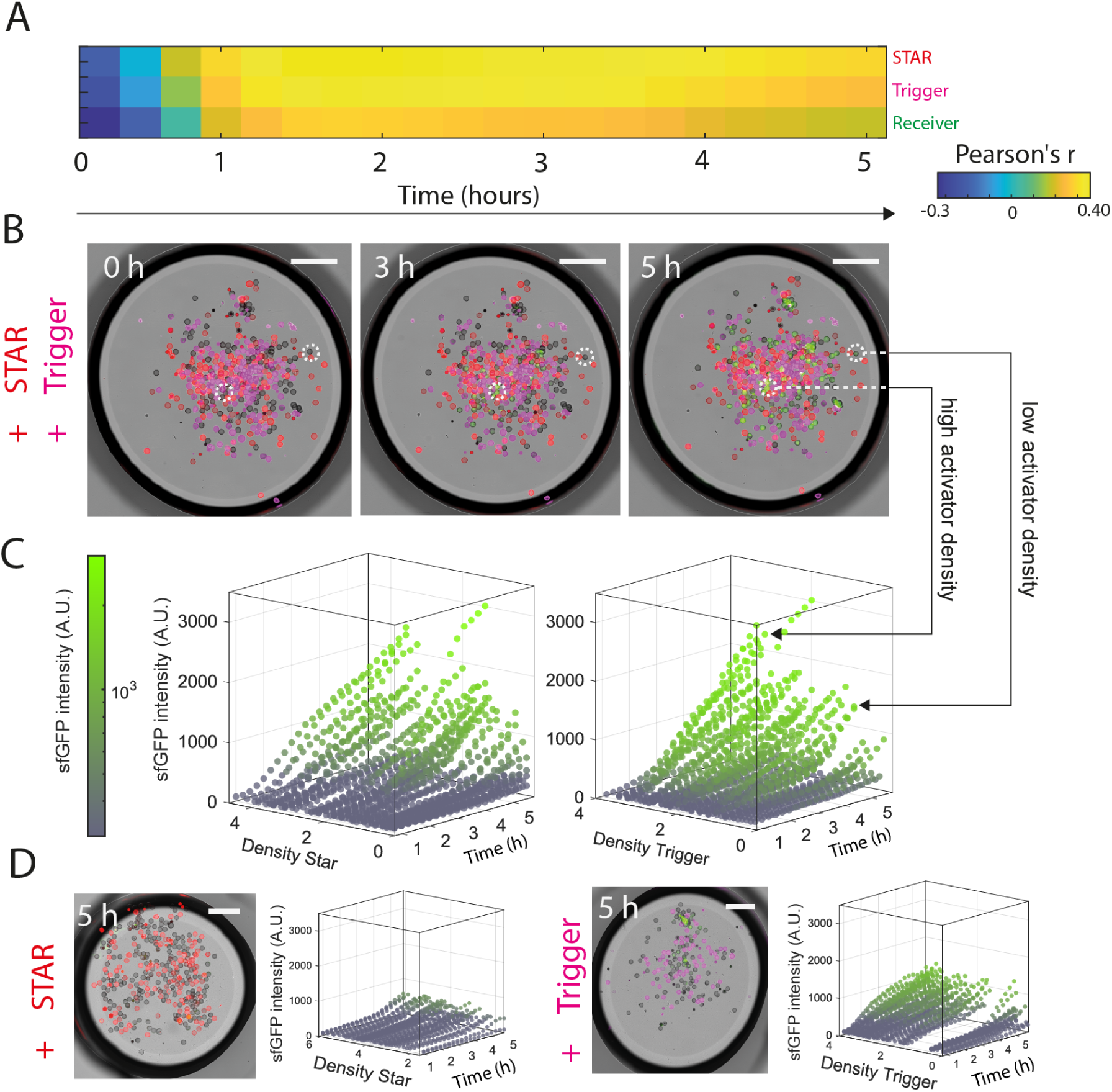
Kinetics of RNA-based communication and AND-gate circuit function in cell mimics. (A) Changes in correlation between receiver fluorescence and the density of each cell mimic type within a radius of 600 µm around the receiver, over time (n = 349 receivers). Heatmap shows correlation coefficients over 5 hours of cell-free expression, averaged across the 5 timelapse samples. (B) Example images from a representative time-lapse experiment containing both STAR and trigger senders. Merge of brightfield and fluorescence channels for sfGFP (green), STAR senders (magenta) and trigger senders (red). (C) Kinetics of sfGFP fluorescence in individual receiver cell mimics. Traces of 70 cell-mimics from 3 selected, representative samples, ordered by local sender density. (D) Kinetic sfGFP intensity traces of individual receivers in control experiments (from 5 samples each per control conditions) that omitted one of the sender types. Scale bars: 450 µm. Density values are the number of cell mimics per mm^2^.

### Spatial arrangement of sender and receiver cell mimics visualizes signal propagation

Analyses of the kinetics and local environments in RNA-based communication between cell mimics revealed spatiotemporal gradients in AND-gate activation in receiver cell mimics. Fold-activation and local effects were most prominent, when the synthesis of both small activating RNAs was high nearby and before diffusion homogenized responses. In manually arranged experiments of clustered sender populations and disperse receiver cell mimics, we observed a clear activation bias closer to trigger clusters with nearby STAR senders, and gradients of activation in more distant receivers (**Fig. S9**). The stronger activation near trigger clusters is likely due to a combination of factors, including slower trigger diffusion, its strong effect on translation activation and leakiness of the STAR-regulated transcriptional terminator. For maximal activation, we prepared senders that produced both small activating RNAs, STAR and trigger, in one sender cell mimic type. To visually demonstrate a spatiotemporal signalling gradient in our system, we prepared a non-random arrangement of communicating cell mimics, where we surrounded a central patch of senders with receivers. To accommodate the spatial arrangement of cell mimics and visualize signal propagation in space and time, we increased the TXTL reagent volume (20 µl) of the sample droplet as compared to the previous experiments, so that the distance of the center with senders was approximately 2 mm to the furthest receivers at the edge. Over the course of the 5-hour reaction, we observed sequential activation of the receivers starting in the center, near the senders (**Fig. 5A**). A kymograph that shows the average sfGFP intensities of 15 receivers at increasing distances from the center, visualizes spatiotemporal signal propagation (**Fig. 5B**). Receivers located closer to the center (<0.8 mm) were activated first, between 1.5 and 2 h, followed by a sequential activation of the more distant receivers until the experiment ended. The kymograph data support the observations from kinetic measurements in randomly distributed samples (**Fig. 4**), indicating that lower sender densities — interpreted here as increased distances from the senders — resulted in delayed sfGFP accumulation in the receivers. Intensities and fold-activation of receivers near the senders were higher than in experiments with random arrangements and two sender types, corroborating our earlier findings that high concentrations of both small RNA signals improve activation and the signal to noise ratio.

**Figure 5.**
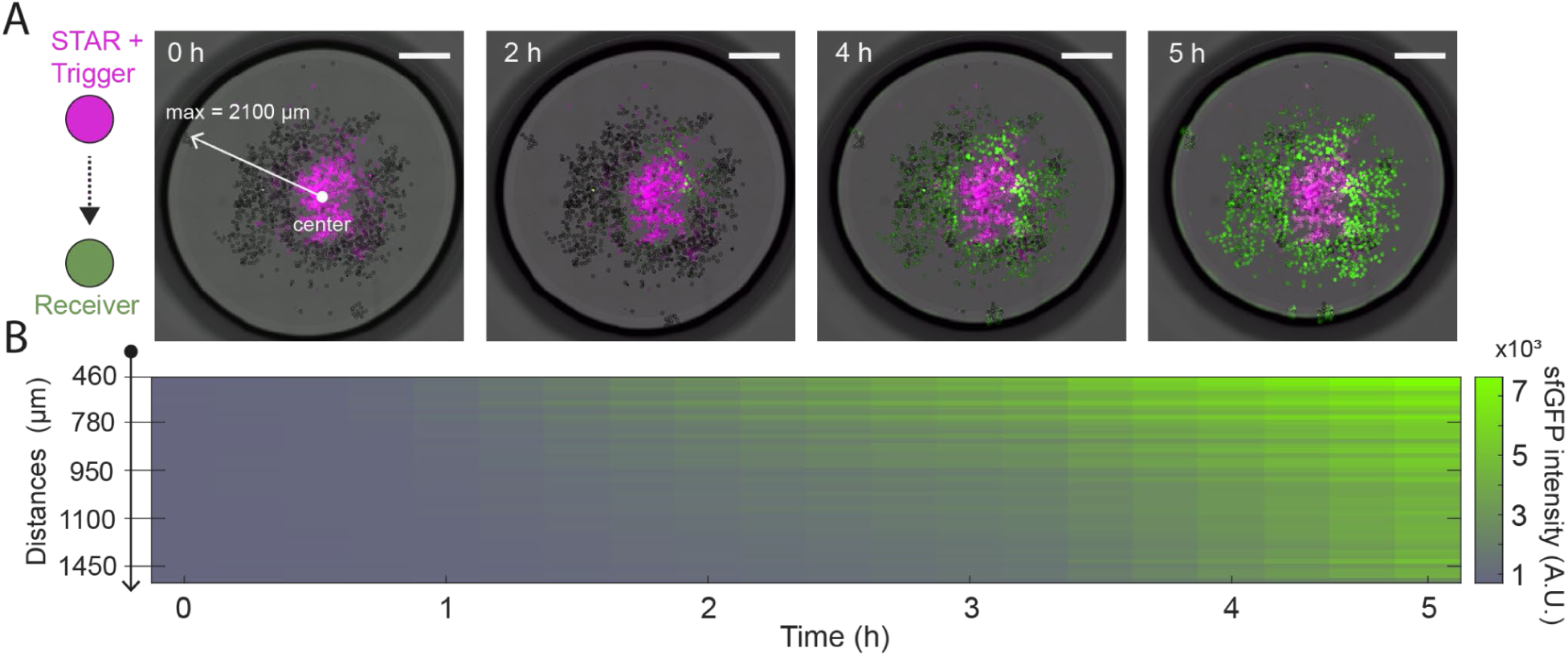
Signal propagation from a centrally localized sender population containing both STAR and trigger plasmids within a single cell mimic type. (A) Time-lapse images of sequential receiver activation. Merge of brightfield and fluorescence channels for sfGFP (green) and STAR+Trigger senders (magenta). The distance from the center of the droplet to the farthest receiver is 2100 µm. (B) Kymograph of the sfGFP fluorescence averaged over 15 receivers along the distance from the closest to the farthest receiver within the droplet. Scale bars: 900 µm

## Conclusions

Our study demonstrates that small, synthetic regulatory RNAs can serve as signaling molecules between synthetic cells, to directly activate transcription and translation in synthetic cells that receive the signals. Specifically, we show activation of an AND gate circuit that is regulated by STAR and trigger RNA signals from two separate sender populations. Using RNA as a signal opens up a vast design space for orthogonal communication channels, as de novo STAR and trigger RNA-regulated switches can be easily engineered via programmable base pairing interactions. ^35,42^ A large repertoire of characterized switches is available, and their performance continues to be improved. ^35,36,39,40,43^ Besides facilitating rational design, small synthetic RNAs are attractive regulators in synthetic circuits because they can act faster than proteins and require less resources for their synthesis. The programmability of cell mimics means that the system could be adapted to other riboregulators and ligand-responsive riboswitches, as well as more complex computations in the signaling circuits. ^42,44,45^

Comparing the effects of the STAR and the trigger on AND gate activation revealed that regulation of the toehold switch by the trigger RNA at the level of translation was tighter than transcriptional termination activated by the STAR regulator, which displayed leaky expression. This behavior was expected based on previous ^37,39,40^ and our own analyses in bulk reactions. Spatially separating the sources for the input signals from the regulated logic gate, enabled us to analyze a distributed logic computation and to compare the ranges of both signaling molecules. Globally, across a receiver population in a sample, the distributed circuit decreased average activation signals and fold-changes. Locally, however, activation was strong in high cell mimic density regions. At short distances between individual cell mimics, we found a higher correlation between trigger sender density and receiver fluorescence than for STAR senders. We hypothesize that this finding can be explained by the potency of trigger-mediated activation of translation and its predicted slower diffusion compared to STAR RNA. Detailed analysis of activation kinetics revealed spatiotemporal gradients of RNA signals that displayed the highest difference between specific activation and undesired sfGFP signals at intermediate time scales.

In the distributed AND gate, leaky expression in the trigger-only condition and high trigger density regions likely arises from a combination of partial target terminator readthrough and the intrinsic activity of the strong promoter in the reporter construct. ^40^ Future optimization could include: (i) incorporating multiple copies of efficient intrinsic terminators to reduce unintended transcriptional readthrough; ^35,39^ (ii) adjusting the relative expression levels of STAR and trigger RNAs by tuning the number of cell mimic senders in the system (e.g. less than 2.5 trigger senders per mm^2^) to balance activation efficiency and minimize background expression; and (iii) integrating additional post-transcriptional control modules, such as designed small RNAs or antisense RNAs, to suppress basal transcriptional leakage and enhance circuit robustness. ^44,46^

In addition to integrating signals from two sources, AND gates are another strategy to effectively reduce leaky expression in the absence of any activator. ^47,48^ If leaky expression is problematic for engineering a synthetic function, combining two slightly leaky regulators in an AND gate, can produce a tight off state in the absence of both signals, while allowing activation only in the presence of both activators. Using senders of both STAR and trigger signals, we demonstrate strong specific activation in receivers located in close proximity to the source of both activators.

In summary, RNA signals will be a useful addition to the communication toolbox for synthetic cell engineering. Transcription of small RNAs is fast and conserves resources. Many orthogonal signals are possible and can be combined for tight regulation. ^49^ For engineering collective responses or spatiotemporal gene expression patterns in communities of synthetic cells, ^16,26,30,50^ RNA-based communication is particularly interesting. Taking advantage of the short lifetime of RNA in cell-free expression reactions, RNA signals will allow the implementation of short-range communication, as opposed to stable, slow-diffusing protein signals and fast-diffusing small molecules.

## Methods

### DNA construction

We based our constructs on an earlier study that implemented an RNA-regulated AND gate in bulk cell free expression reactions. ^37^ The activator constructs, J23119-Trigger3-T500 and J23119-STAR6-T500 were used without modification. The plasmids J23119-S6T3-sfGFP-T500, J23119-S6-sfGFP-T500 and J23119-T3-sfGFP-T500 were modified to include TetR, resulting in the fusion protein TetR-sfGFP. Golden gate was performed using the NEBuilder HiFi DNA Assembly (NEB) and NEBuilder Assembly Tool for primer design. Plasmids were purified using Midiprep kits (NucleoBond®Xtra, Macherey-Nagel) and resuspended in UltraPure DNase/RNase-Free Distilled Water (Thermofisher).

### TXTL extract preparation

TXTL extract preparation followed a previously described protocol. ^51^ *E. coli* BL21 Rosetta 2 were streaked overnight on an agar plate containing chloramphenicol. One colony was picked and inoculated overnight in 50 mL 2xYT supplemented with chloramphenicol for growth at 37°C. After a minimum of 15 hours, 20 mL of the stationary culture was used to inoculate 400 mL of 2xYT + P media (16 g/L tryptone, 10 g/L yeast extract, 5 g/L sodium chloride, 7 g/L potassium phosphate dibasic, 3 g/L potassium phosphate monobasic) in a 1 L baffled flask. Cells were grown at 40 °C and 200 RPM to 3.0 ± 0.2 OD_600_. Centrifuge bottles were filled up to 300 mL and centrifuged for 10 minutes at 4,000xg at 4 °C and supernatants were discarded. The pellets were washed three times with 25 mL buffer S30A (50 mM Tris-base, 14 mM Mg-glutamate, 60 mM K-glutamate, 2 mM DTT, brought to pH 7.7 with acetic acid). The washing steps were followed by a centrifugation step at 4,000xg at 4 °C for 10 minutes. A fourth centrifugation step at 3,000xg at 4 °C for 10 minutes enabled the removal of the remaining traces of the buffer. The pellets were then resuspended in 1 mL of buffer S30A per gram of pellet and supplemented with 0.5 mg/mL of lysozyme (from chicken egg, >40,000 units/mg, Sigma). The resuspended pellets were incubated for 10 minutes on ice. 1 mL of the suspension was aliquoted into 1.5 mL reaction tubes. The pellet suspensions were then lysed with a sonicator (QSonica Q125 with a 3.175 mm diameter probe, 50% amplitude, 20 kHz, and 10 seconds ON/OFF pulses). Each sample was sonicated until reaching 250 J input. Using a 100 mM stock solution, 1 mM of DTT was added to each crude lysate immediately after sonication. The cell lysate was centrifuged for 10 minutes at 4 °C and 12,000xg. The supernatant was removed and incubated at 37 °C under shaking at 200 RPM for 80 minutes. After the run-off reaction, the supernatant was centrifuged for 10 minutes at 4 °C and 12,000xg. The supernatant from the centrifugation step was dialyzed for 3 hours against buffer S30B (50 mM Tris-base, 14 mM Mg-glutamate, 60 mM K-glutamate, 2 mM DTT, pH 8.2) in a 10k MWCO cassette (Thermofisher). Finally, the dialyzed extract was centrifuged for 10 minutes at 4 °C and 12,000xg. The supernatant was aliquoted, snap-frozen into liquid nitrogen, and stored at −80 °C.

### TXTL reaction

The final TXTL reaction mixture is composed of the following reagents: 33% v/v of *E. coli* extract, 10 mM ammonium glutamate; 1.2 mM ATP; 0.850 mM each of GTP, UTP, and CTP; 0.034 mg/mL folinic acid; 0.175 mg/mL yeast tRNA; 2 mM amino acids; 30 mM 3-PGA; 0.33 mM NAD; 0.27 mM CoA; 1 mM putrescine; 1.5 mM spermidine; 57 mM HEPES, 2.0 % PEG 8000; 10 mM Mg-glutamate; 130 mM K-glutamate. For the plate-reader experiment, plasmid DNA concentrations were set to 10 nM for the STAR and trigger plasmids and 2 nM for the AND gate reporter plasmid.

### Cell mimic production

Cell mimics were produced according to previously published protocols with adjusted polymer composition. ^31^ Briefly, we prepared flow focusing, microfluidic chips by partially coating them with PVA solution and baking at 120°C overnight before storing them at room temperature until use. To create the double emulsions, we used an aqueous outer phase of 2.5% (w/v) poloxamer P 188, an aqueous inner phase of 63% (v/) glycerol and 2 % (w/v) poloxamer P 188, and an organic middle phase composed of 15 mg 2,2 dimethoxy 2-phenylacetophenone dissolved in 350 μl 1-decanol, 325 μl glycidyl methacrylate, 325 μl trimethylolpropane triacrylate and 10 μl Span 80. Each phase was pumped into the chip using syringe pumps and the flow rate adjusted to create a stable stream of double emulsion.

The double emulsion was collected in 2 mL tubes and further diluted 1:1 with outer phase. The emulsion was then spread out on a weighing boat and polymerized for 30 s with 365 nm UV light at 200 mW/cm^2^. After polymerization, we added ethanol to the suspension to reach a 70% (v/v) solution and transferred the cell-mimics to a fresh tube for storage. Cell-mimics were washed with pure ethanol before pegylation with an aqueous, 50% (v/v) ethanol, 500mM methoxy-PEG-amine (MW 750) solution at 37°C overnight. To stain cell-mimics, we added 0.5mM amine conjugated fluorescent dye (CF™ 633 amine #SCJ4600035 or CF™ 555 amine #SCJ4600019 from Sigma-Aldrich) to the pegylation solution, which covalently linked to the cell-mimics.

### Cell mimic DNA loading

DNA was immobilized by using its high binding affinity to synthetic clay hydrogels physically trapped within the cell mimics. To load cell-mimics with DNA, they were first thoroughly washed with water (5 times), pelleting them using a tabletop minicentrifuge. Then the clay pre-hydrogel was diffused into the cell-mimics by incubating them overnight in a 1.6% suspension of synthetic clay Laponite® XLG (BYK Additives). To remove external clay, the cell-mimics were washed once with water. Right after washing with water, cell mimics were incubated with DNA solutions (150 nM AND gate plasmid and 50 nM PRS316-240xtetO ^52^ (Addgene #44754) for receivers and 750 nM STAR and/or trigger plasmid for activators in water) for five minutes. Finally, the clay-DNA mix within cell mimics was gelled by adding an excess of 100 mM HEPES pH 8 solution and incubating for 10 minutes. After gelling, the cell-mimics were stored in 70% EtOH 100 mM HEPES solution, in the freezer, until use. This procedure condenses the clay-DNA hydrogel into a nucleus-like structure that is smaller than the surrounding polymer shell. ^31^

DNA loading was quantified by comparing cell mimics, loaded with fluorescently labeled linear DNA, with water in oil emulsion droplets with similar size and corresponding DNA loading concentrations. Fluorescent DNA was created by PCR amplification of a 1kb fragment using a TYE 665 labeled primer (IDT), creating DNA templates with one fluorophor per amplicon. To create a calibration curve, 750, 500, 100 and 0 nM labeled DNA in water was emulsified in light mineral oil containing 1% Span80 surfactant by gentle racking of the tube. A large quantity of polydisperse droplets were imaged for every DNA concentration. Of those droplets, the fluorescence data was extracted for droplets in the size range of cell mimics (diameter 60 - 80 μm) using the thresholding and analyze particle functions in Fiji/ImageJ. Fluorescence data for cell mimics loaded with different amounts of fluorescently labeled DNA suspended in a 100 mM HEPES buffer was recorded with the same microscope settings. Cell mimic fluorescence was equally extracted using Fiji/ImageJ. To calculate the calibration curve, a MATLAB script was used computing the slope and intercept of a linear regression model corresponding to the droplet fluorescence (**Fig. S5A**). From this calibration curve, the actual DNA loading, for individual cell mimics, was calculated and compared with the loading input concentrations (Fig. S5 B). Using the experimentally determined DNA loads of senders and receivers, we calculated the approximate total DNA concentrations in a 5 µl sample containing 100 cell mimic spheres with a 70 µm diameter.

### Cell mimic transcription and translation reactions

A typical cell-mimic reaction consisted of 1 µl concentrated cell-mimics, suspended in 100mM HEPES pH8, in a total TXTL reaction of 5 µl. 3 µl of the suspension were spotted into a 35 mm Lumox® dish (Sarstedt), with a vacuum grease rim, and covered with a cover glass sealing the grease. The membrane of the Lumox dish allows for gas exchange while the grease and cover glass function as spacer and prevent evaporation. Up to 5 sample droplets could be spotted into one Lumox dish. Samples were incubated at 29°C for 5h and then imaged. Larger experiments, with spatially arranged cell-mimic populations, were sequentially placed into the Lumox® dish, first spotting the receiver cell mimics, followed by sender cell mimics in the center which displaced the receivers into a surrounding ring, and then covering them with TXTL reagents. Endpoint and time-lapse imaging was performed using a Nikon Eclipse Ti2-E inverted microscope with a CFI S Plan Fluor ELWD 20x/0.45 (MRH08230) objective and a pco.edge 4.2 bi camera. CF 633 labeled cell-mimics were imaged in the mCherry channel and CF 555 labeled cell-mimics in the Cy5 channel. For time-lapse series, images were recorded every 15 minutes. Large image acquisition was done using a custom JOBS protocol to automatically image and stitch individual tiles.

To reduce variability, all experiments were performed within a single month using the same batch of TXTL reagents and the same batch of cell mimics loaded with the different genetic constructs. Microscopy was carried out under identical imaging conditions for all experiments. For each experimental setting (combinations of cell mimic populations), data collection was distributed across two experimental days to ensure reproducibility. On each day, the numbers of cell mimics per sample were varied and spotted into Lumox® dishes, which each contained up to four samples. For example, in the endpoint experiments involving the population containing the AND gate together with both activators, a total of 8 Lumox® dishes, each containing three or four samples, were prepared on two separate days (for an overview of all samples see **Fig. 2A**).

### Image Analysis Pipeline

After image acquisition, the image sizes were reduced four-fold using ImageJ before analysis. Images were processed with a custom script in MATLAB. Briefly, cell mimics were segmented in the bright-field channel using the built-in Circular Hough Transform function, followed by further analysis and filtering with functions from the Image Processing Toolbox. The segmented cell mimics were then classified as either receivers or senders based on intensity thresholding in the Cy5 and mCherry fluorescence channels, using manually determined thresholds. The sfGFP channel was used to extract fluorescence intensity from receiver cell mimics. Euclidean distances between each receiver and all senders were calculated. These distances were then used to count the number of sender and receiver mimics within a specified radius around each receiver. The local density of cell mimics was computed using the formula *n*/*A*, where *n* is the number of cell mimics of a given type within the area A, defined as the area of a circle with radius *R* around the receiver. Finally, the Pearson’s r correlation coefficient was calculated for each sample by correlating the fluorescence intensity of each receiver with the local density of each cell mimic type surrounding it. The final Pearson’s r value presented in Fig. 3 represents the average of all correlation coefficients computed across individual samples. For the analysis of activation near spatially arranged sender clusters, receiver cell mimics were manually selected in the bright field channel along a line interpolating the two sender clusters. Fluorescence values were extracted from the sfGFP channel for circular regions corresponding to the selected cell mimics.

### RNA property prediction

RNA signal secondary structures were predicted with ViennaRNAfold ^53^ and fed into RNAComposer to predict their tertiary structures. ^54^ To estimate the diffusion coefficients, HullRad was used with the 3D structures as input. ^55^

## Author contributions

H.N., F.X.L. and I.B. conceived, planned, and designed the study. F.X.L., I.B., E.B. and T.M. performed the experiments. F.X.L. performed the data analyses. F.X.L. and H.N. wrote the manuscript with the input of all authors. H.N. acquired funding and supervised the work. All authors approved the final manuscript.

## Notes

The authors declare no competing financial interest.

## Acknowledgements

This work was supported by the Deutsche Forschungsgemeinschaft (DFG, German Research Foundation) grant NI 2040/1-1.

## Supporting Information

Figure S1: Correlations between cell mimic types and average receiver sfGFP intensity in experimental samples.

Figure S2: Spatial coordinates do not influence the sfGFP production of the receiver cell mimics.

Figure S3. Receiver fluorescence in all samples including the extended range of trigger sender cell mimics in control experiments.

Figure S4: Influence of sender cell mimic counts on Pearson correlation coefficients.

Figure S5: Quantification of cell mimic DNA loading.

Figure S6: Structure and diffusion coefficient prediction for signal RNAs.

Figure S7: Sources of leaky expression.

Figure S8: Kinetics of RNA-based communication and AND-gate circuit function in cell mimics color-coded according to the receiver density.

Figure S9: Spatially arranged sender clusters and their influence on local AND gate activation.

Movie 1: Movie of the rotating 3D plot of individual receiver sfGFP fluorescence intensities across all experiments as shown in Figure 3D.

Movie 2: Selected time-lapse movie of a sample containing both STAR (red) and trigger (magenta) senders. sfGFP fluorescence in receivers is shown in green.

