## Supporting Information for "RNA-based communication in heterogeneous populations of cell mimics"

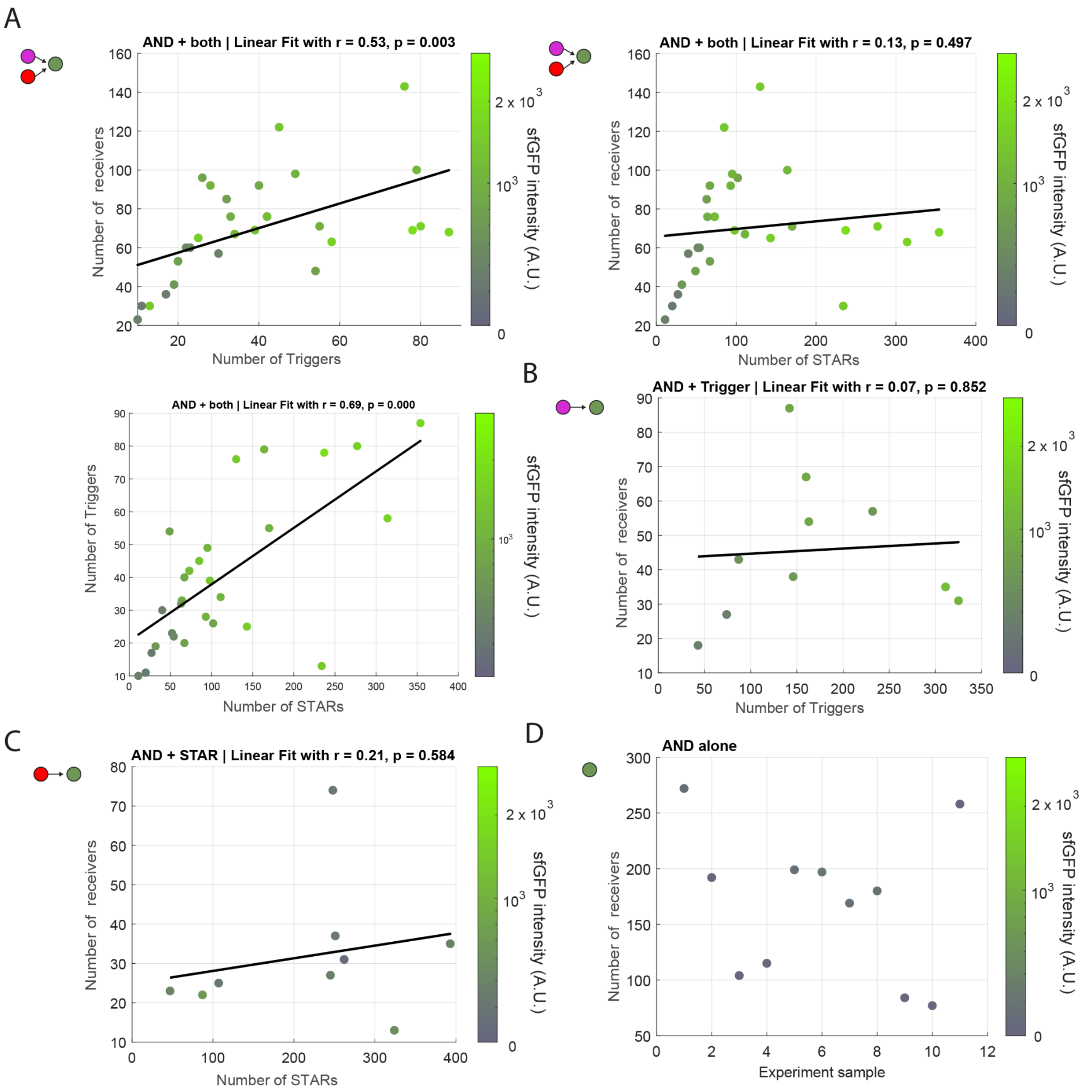


**Supplementary Figure S1.** Correlations between cell mimic types and average receiver sfGFP intensity in experimental samples. For: A) the AND gate in presence of the two types of sender cell mimics;  B) the AND-gate with only trigger senders; C) the AND-gate with only STAR senders; D) the AND-gate without senders. Pearson correlation coefficient (r) and p-value (p) are displayed above each plot (except for receivers with the AND-gate alone).

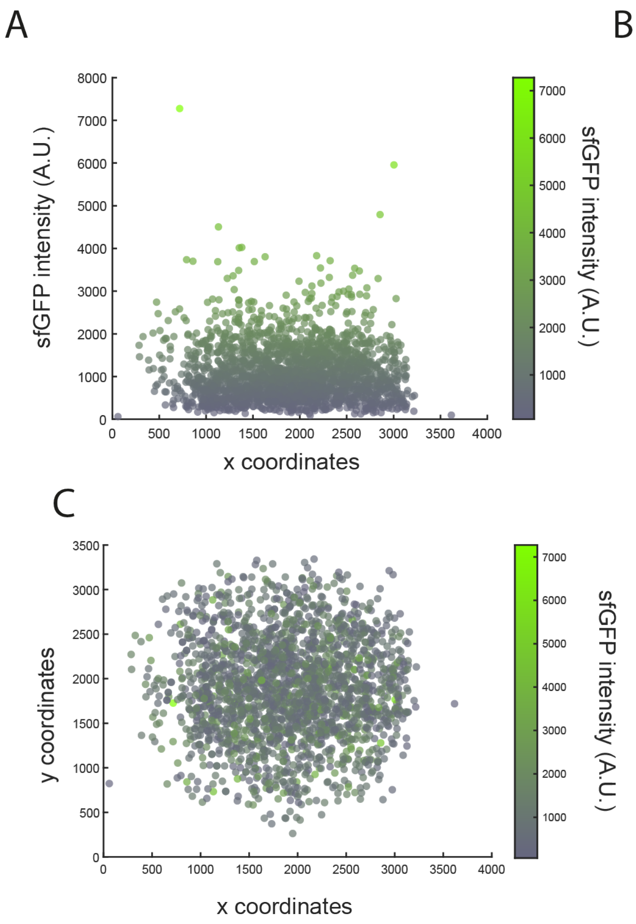


**Supplementary Figure S2.** Spatial coordinates do not influence the sfGFP production of the receiver cell mimics. A-C) Influence of x-y coordinates on sfGFP intensity. Data points show the coordinates and sfGFP intensities of 2037 individual receiver cell mimics from 29 samples containing the full AND gate (STAR and trigger senders plus receivers). Peason correlation coefficients indicated no correlation between sfGFP intensity and x-y coordinates (Pearson's r < 0.1).

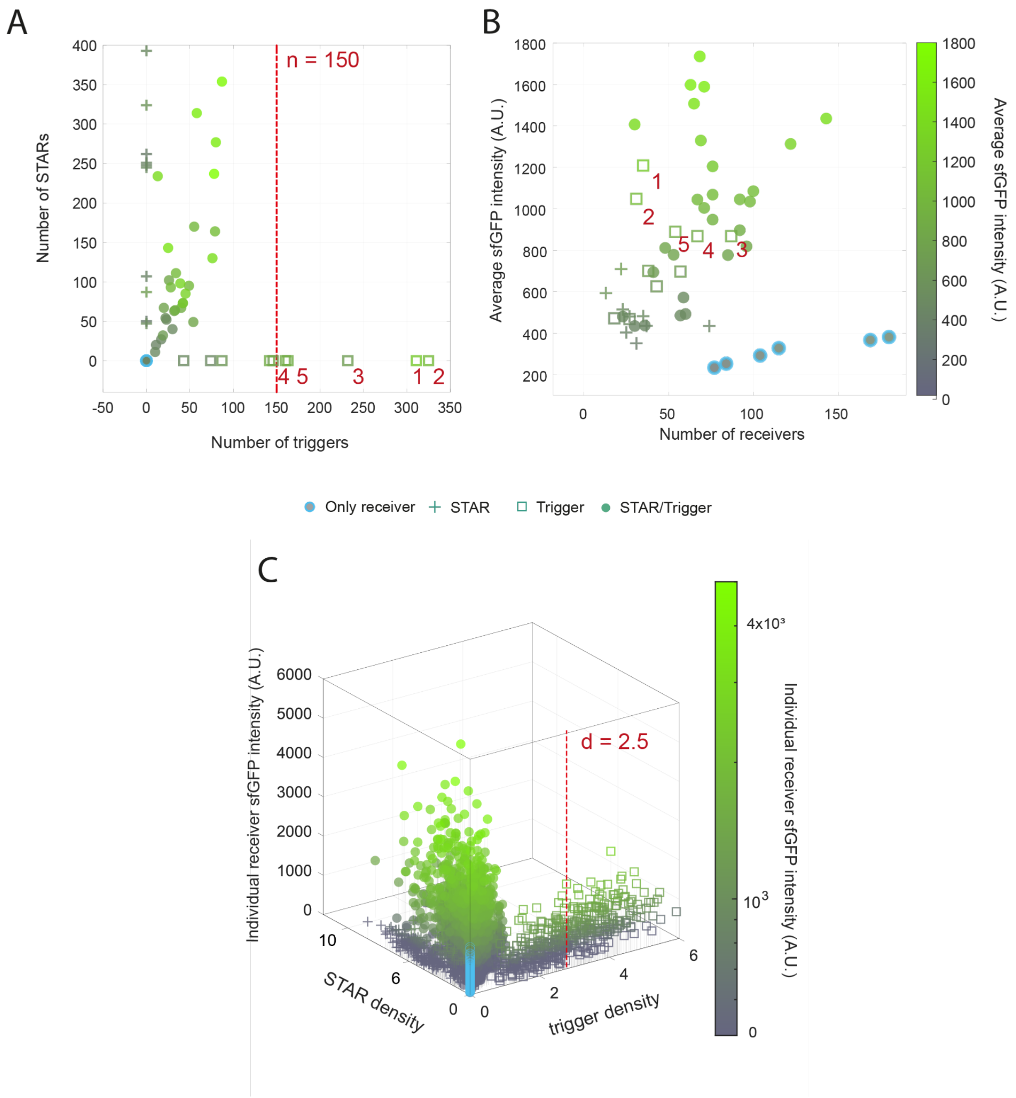


**Supplementary Figure S3.** Receiver fluorescence in all samples including the extended range of trigger sender cell mimics in control experiments. A) Analysis of sfGFP endpoint fluorescence values averaged across all receivers in a sample as a function of trigger and STAR counts. B) Analysis of sfGFP endpoint fluorescence values averaged across all receivers in a sample as a function of receiver counts. Red numbers indicate the samples from the extended trigger range in (A). The extended range for the trigger count starts at 150 (dashed line). C) sfGFP fluorescence values of each individual receiver across all experiments, including controls that omitted a sender type, plotted against local sender densities in a radius of 600 µm around each receiver. Density values indicate the number of cell mimics within the radius around the receiver divided by the circle area. Density values are the number of cell mimics per mm$^2$. The extended range for trigger density starts at 2.5 (dashed line).

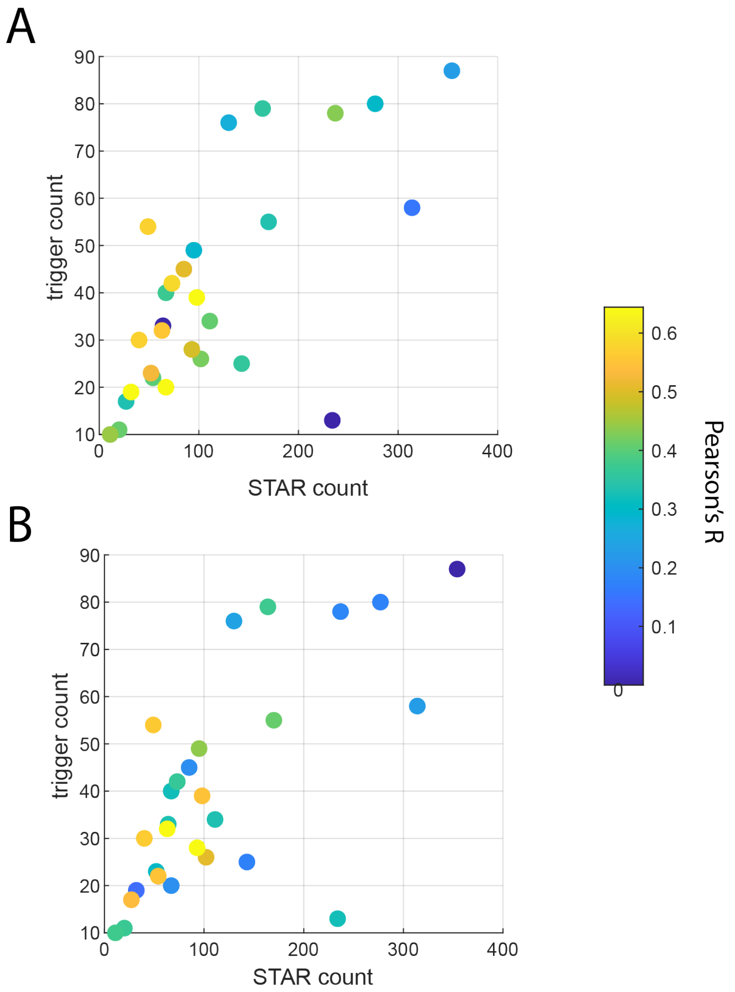


**Supplementary Figure S4.** Influence of sender cell mimic counts on Pearson correlation coefficients. A) Pearson correlation coefficients with the trigger senders (r, color code) in respect to STAR and Trigger sender counts in all experimental samples. B) Pearson correlation coefficients with the STAR senders (r, color code) in respect to STAR and Trigger sender counts in all experimental samples. Pearson's r correlations are averaged over all correlations with the same sender type present within a given sample for a radius of 600 µm.

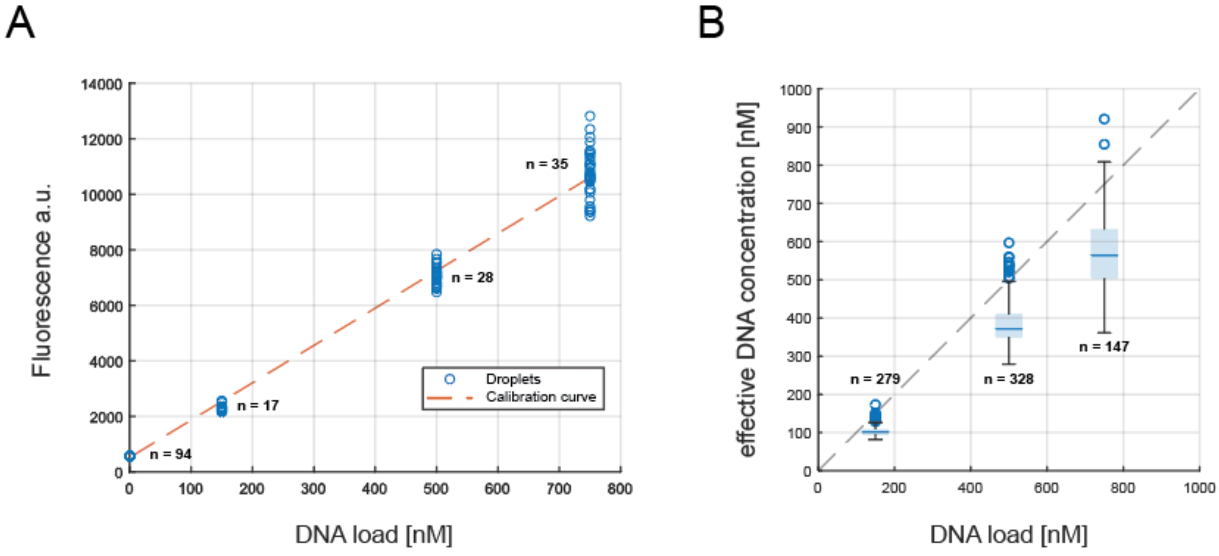


**Supplementary Figure S5.** Quantification of cell mimic DNA loading. A) DNA loading calibration curve in emulsion droplets (orange dashed line). We emulsified fluorescently labeled DNA at specific concentrations in mineral oil and measured the fluorescence of droplets in the size range of cell mimics with diameters between 60-80 µm (blue circles, n = number of droplets per concentration). The calibration curve was calculated as simple linear regression with a y-intercept at the background fluorescence for droplets without DNA. B) Comparison of effective DNA immobilized in individual cell mimics (y axis) versus DNA concentration used for cell mimic loading (x axis), calculated with the calibration shown in (A). Box plots show three loading conditions of 150, 500 and 750 nM DNA with calculated effective DNA immobilized within the cell mimics. Blue lines show average DNA loading while the box marks the upper 0.75 and lower 0.25 quantile (blue circles are outliers, n = number of cell mimics per concentration). The grey, dashed line marks the ideal 1:1 loading:immobilization.

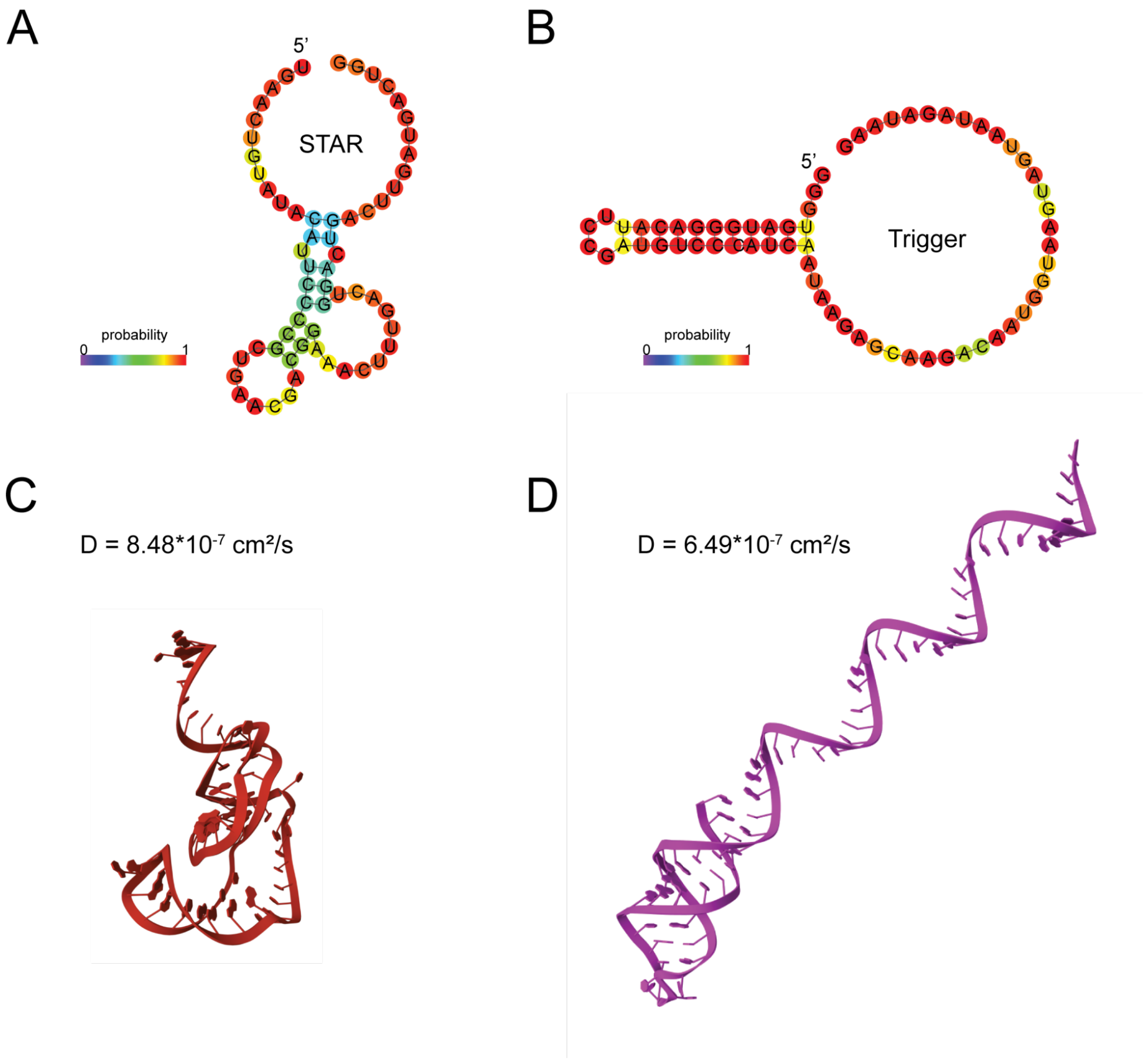


**Supplementary Figure S6.** Structure and diffusion coefficient prediction for signal RNAs. A) Secondary structure of STAR calculated using ViennaRNAfold. The colorbar indicates the base-pair probabilities. B) Secondary structure of Trigger calculated by ViennaRNAfold. C) Tertiary structure prediction of STAR by RNAcomposer and its diffusion coefficient calculated by HullRad. D) Tertiary structure prediction of Trigger by RNAcomposer and its diffusion coefficient calculated by HullRad. See methods for references to the prediction tools.

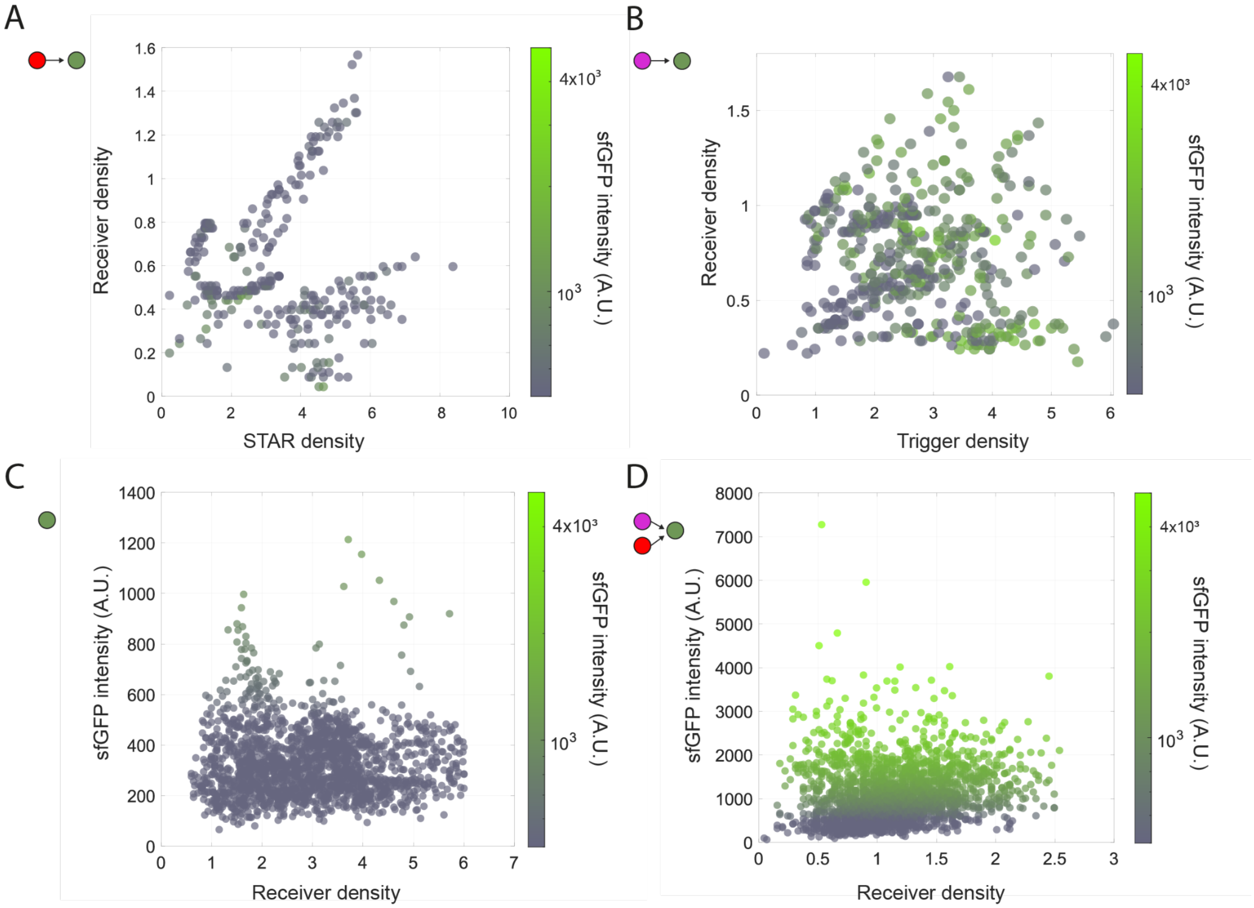


**Supplementary Figure S7.** Sources of leaky expression. Individual receiver sfGFP intensities for different experimental configurations. A) Receivers and STAR senders only; B) Receivers and trigger senders only (including the extended range for trigger counts >150); C) Receivers alone; D) Receivers with both STAR and trigger senders. Density values indicate the number of cell mimics within the radius around the receiver divided by the circle area. Density values are the number of cell mimics per mm$^2$ analyzed in a 600 μm radius around each receiver.

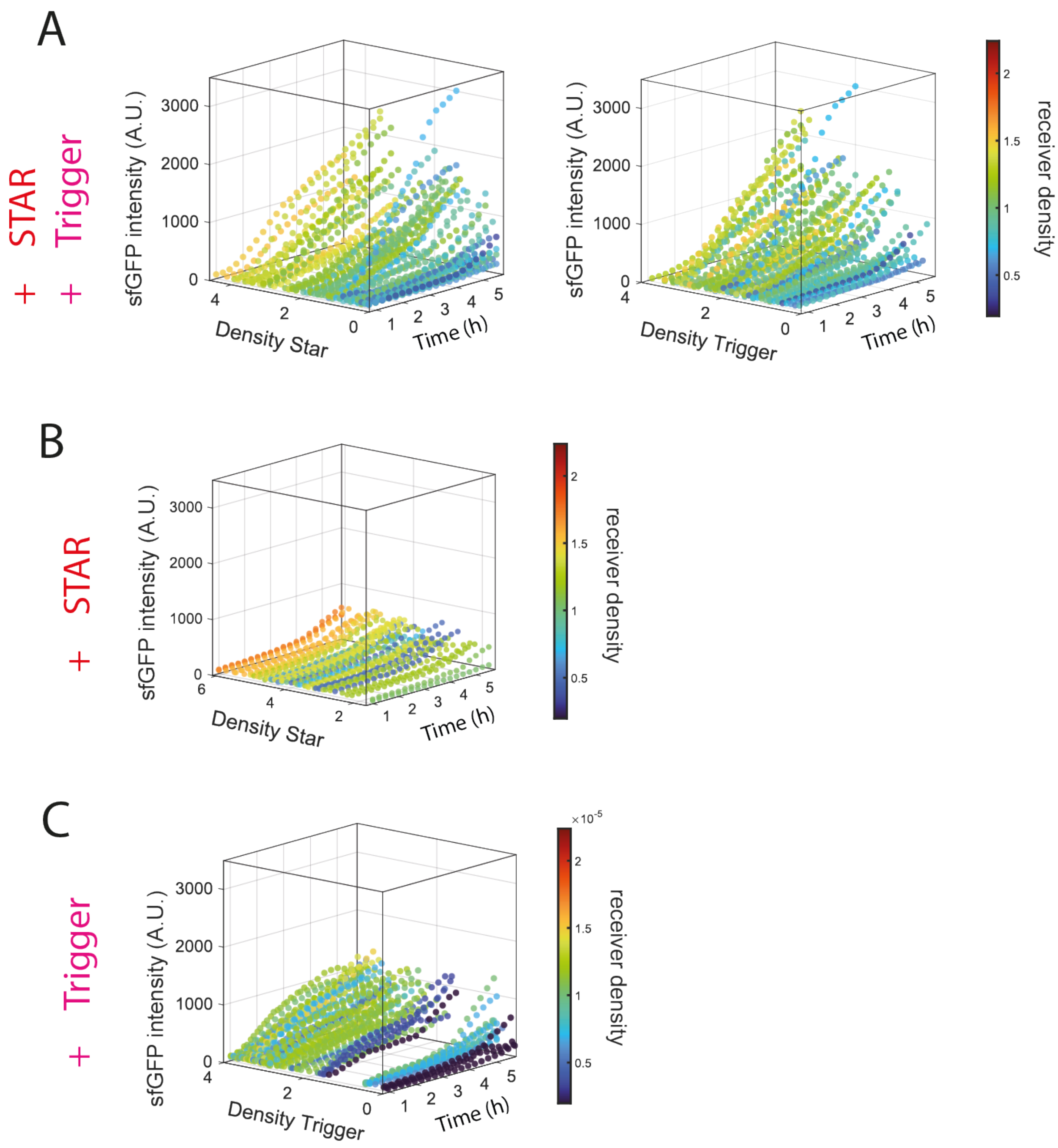


**Supplementary Figure S8.** Kinetics of RNA-based communication and AND-gate circuit function in cell mimics color-coded according to the receiver density within a radius of 600 µm around the receiver. (A) Kinetics of sfGFP fluorescence in individual receiver cell mimics. Traces of 70 cell-mimics from 3 selected, representative samples, ordered by local sender density. (B) and (C) Kinetic sfGFP intensity traces of individual receivers in control experiments (from 5 samples each per control conditions) that omitted one of the sender types. Density values are the number of cell mimics per mm$^2$.

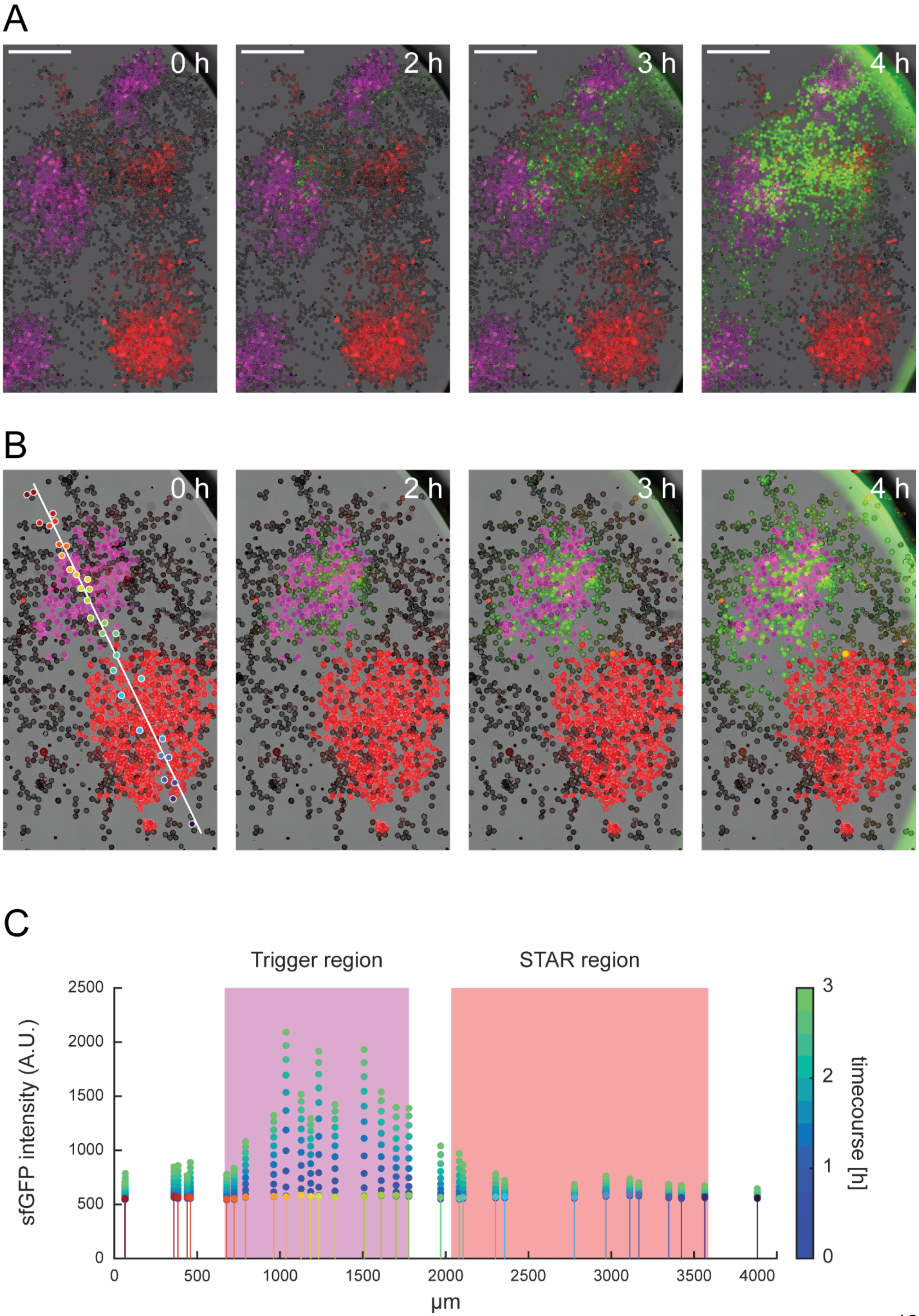

A
0 h
2 h
3 h
4 h
B
0 h
2 h
3 h
4 h
C
Trigger region
STAR region
sfGFP intensity (A.U.)
2500
2000
1500
1000
500
0
0
500
1000
1500
2000
2500
3000
3500
4000
µm
timecourse [h]
3
2
1
0

**Supplementary Figure S9.** Spatially arranged sender clusters and their influence on local AND gate activation. A) Merged-channel, timelapse images of gate activation. The sample contained multiple clusters of trigger (magenta) and STAR (red) sender cell mimics with dispersed receivers (grey). Gate activation is strongest at the intersection of multiple, neighboring STAR-trigger clusters with a bias towards trigger senders. Scale bar is 1 mm. B) Merged-channel, timelapse images of gate activation. Two distinct clusters of trigger (magenta) and STAR (red) sender cell mimics with interspersed receivers (grey). The white diagonal, in the 0h-panel, measures 4 mm and marks a virtual line from one center of a cluster to the other. The colored points mark selected receiver cell mimics used for data analysis along the virtual line. C) Time course of fluorescence signals from selected cell mimics along the line in panel B. The color of the circular markers at 0h corresponds to the points in panel B. Magenta and red background demarcate the sender clusters on the line. Fluorescence is strongest close to and within the trigger region, with an offset towards the STAR region, indicating the local AND gate activation as well as leaky expression as expected from results with random cell mimic distributions.
